# A tailored variant filtering procedure for multi-breed and multi-species unbalanced animal SNP collections

**DOI:** 10.1101/2025.10.14.682050

**Authors:** Barbara Lazzari, Marco Milanesi, Andrea Talenti, Arianna Bionda, Yefang Li, Lin Jiang, Johannes A. Lenstra, Philippe Bardou, Gwenola Tosser Klopp, Paola Crepaldi, Licia Colli, The VarGoats Consortium

## Abstract

Technological advancements and decreasing costs of whole-genome sequencing have generated a huge amount of resequencing data. Large-sized datasets, spanning several species and/or populations can now be assembled easily. However, they vary widely in geographical provenance, and sample sizes, with taxonomic groups varying from single to hundreds of entries. Consequently, standard filtering approaches may bias the representation of groups or gene pools. Commonly adopted variant filters relying on minor allele frequency (MAF) and linkage disequilibrium (LD) are not adequate because LD patterns and allele frequencies differ substantially within datasets representing both local and global diversity of multiple populations and species. Thus, by using the VarGoats 1000 goat genome project, we devised a novel approach which avoids the biases of the standard filtering procedures by adopting within-population subsampling, minor allele count (MAC) and marker spacing (bp-space) as filters. Starting from a quality-filtered dataset of >28M SNPs from 1372 animals, we generated a dataset of <14M markers and 750 individuals, complying with the initial requirements and facilitating further computational steps.

## Introduction

After their introduction about 20 years ago, high-throughput sequencing technologies have been increasingly exploited in genetic studies and have now become a routine tool to characterize genome-wide variation in wild and domestic species. The growing availability of reference genome sequences, together with the decreased costs and the standardized data production, has led to the accumulation of resequencing data in public databases, so that it is now possible to assemble large genomic datasets for several domestic and wild species, such as cattle^1^, sheep^2^, pigs^3^; buffaloes^4^, donkeys^5^ and canids^6^. For example, the NCBI SRA database (accessed on August 27^th^, 2025) encompassed 98,591 sequence data entries for cattle representing breeds from all over the world, 2,100 sequencing data entries for the cosmopolitan grey wolf, and more than 4,200 genome data for six species of Darwin’s finches.

Populations from different geographical areas or with different evolutionary (e.g. domesticated vs. wild relative species) or demographic (e.g. small vs. large effective population size, bottlenecks, etc.) histories usually differ in terms of allele frequency and linkage disequilibrium (LD)^7–9^. This especially applies to domestic species, where human management has reduced between-breeds gene flow^10,11^. As shown by SNP data in worldwide goats, the number of markers that are found in linkage differs for populations from Africa, Europe and Southwestern Asia^12^. There are also remarkable differences in the distribution of molecular variation within and between continents: Central and Northern European breeds tend to form well defined and separated clusters, while in Southern Europe and within the African continent the goat populations are less differentiated with regional gene pools being shared by several populations of neighboring countries^12^.

Ascertainment bias in SNP panels is commonly caused by over-representation of specific gene pools in the panel of populations used to select the variants^13,14^. Thus, individuals from populations included in the discovery panel will consistently display higher levels of variation than the representatives of other gene pools. This also applies to large-scale WGS datasets assembled from public sources or different projects in which popular breeds or gene pools are often overrepresented. During the filtering steps after the raw sequence data Quality Control (QC) and preceding data analyses, the predominance of one or a few gene pools leads to underrepresentation of the highly informative variation within minor gene pools.

Filtering datasets is of fundamental importance, but different ways of filtering may return completely different results^16,17^. The usual filtering relies on the minor allele frequency and LD-based pruning, i.e. the removal of variants that are in linkage with another variant beyond a specific threshold value overall^18^. In addition, commonly used default program settings for filtering may remove Hardy-Weinberg equilibrium deviations that are important to understand population structure^15^.

Altogether, these approaches may remove low-frequency variants typical of minor gene pools, with the ultimate effect of introducing biases at multiple levels in the filtered dataset. This also affects the estimation of LD, which depends on the allelic frequency of SNP pairs^11^. As a consequence, analysis methods that rely on allelic frequencies can be negatively affected^14^, thus confounding the detection of population structure and the estimation of allele sharing etc., particularly when the minor gene pools are involved.

The VarGoats project represents a remarkable example of a global-scale assemblage of whole-genome sequences deriving from the aggregation of project-specific data from multiple providers (projects VarGoats and NextGen, https://projects.ensembl.org/nextgen/) and publicly available sources (NCBI BioSample, https://www.ncbi.nlm.nih.gov/biosample/). The current VarGoats dataset is an extended version of the one described in Denoyelle *et al*. (2021)^19^ which encompasses 1,372 animals from 132 local and transboundary domestic goat populations, and nine wild goat species collected in Europe, Africa, Asia, and Oceania (https://www.goatgenome.org/vargoats.html; Table 1, Supplementary Table S1). VarGoats WGS data are accompanied by metadata as individual sex, provenance, geographical coordinates of sampling locations (latitude and longitude) and other accessory information, which are particularly useful in evaluating and validating the results of different SNP filtering procedures. However, the VarGoats dataset is not normalized in terms of number of animals per breed and species, which ranges from 1 to 163. This is particularly relevant for the nine wild *Capra* species that together are accounted for by 30 animals overall, which risks to further penalize variants from these underrepresented, overly different individuals in a dataset strongly unbalanced towards *Capra hircus*. All these factors, together with the need to create an easy-to-manage representative dataset, called for the implementation of an *ad hoc* SNP filtering procedure, also applicable to other datasets sharing the same issues.

**Table 1.**
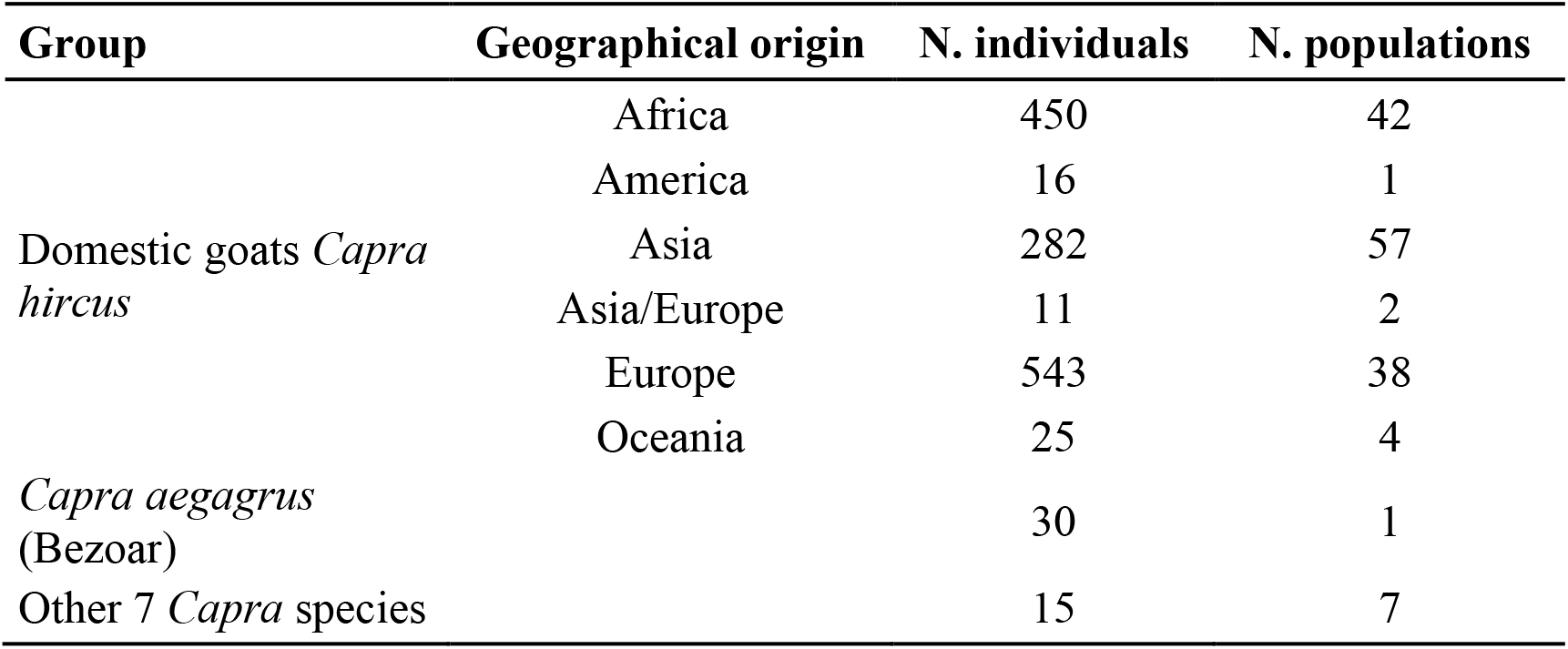
Composition of the VarGoats dataset.

| Group | Geographical origin | N. individuals | N. populations |
| --- | --- | --- | --- |
| Domestic goats *Capra hircus* | Africa | 450 | 42 |
|  | America | 16 | 1 |
|  | Asia | 282 | 57 |
|  | Asia/Europe | 11 | 2 |
|  | Europe | 543 | 38 |
|  | Oceania | 25 | 4 |
| *Capra aegagrus* (Bezoar) |  | 30 | 1 |
| Other 7 *Capra* species |  | 15 | 7 |

## Results and discussion

After mapping to the ARS1 *C. hircus* genome (GCA_001704415.1), quality filtering and phasing, 28,321,956 autosomal biallelic SNPs were retained out of a starting set of 77,280,295 variants. This dataset (referred to as the 28M dataset) represented the starting point for the following procedures.

### Within-breed subsampling for sample size harmonization

We first considered the uneven distribution of animals across breeds and species. After removal of 56 samples that did not reach the minimum quality standards required to be included in further analyses, 13 populations were represented by transboundary breeds sampled in different geographical areas. Because these breeds may be reared under different management conditions in different countries, we decided to split animals based on both breed and country of provenance, thus creating 163 country_breed groups (149 from goat, 2 from bezoars and 12 from other wild goats), comprising 1 to 163 individuals.

A distribution of country_breed sample sizes guided our choice to set the maximum number of individuals per country_breed at eight, to avoid the overrepresentation of breed-specific gene pools. We reduced the size of the 35 country_breed groups with more than eight animals by using the subsampling approach implemented by BITE v2.0^21^ which maximizes the genetic variability of the eight retained individuals and removes the most highly related animals. This subsampling retained 750 out of 1,372 individuals: 720 domestic (DOM) goats in 149 country_breed groups; 16 bezoars (BEZ) in 2 country_breed groups; and 14 other wild goats (WIL), in 12 country_breed groups. The 28,321,956 phased SNPs of the selected 750 samples (referred to as the 28M_BALANCED dataset) were used as the starting point for the subsequent thinning procedure.

### Selection of minor allele threshold value

For an adequate representation of the variation of domestic, bezoars and wild goats, we extracted three overlapping sets of variants from the 28M_BALANCED dataset, including the subsets of SNPs present in the DOM, BEZ and WIL groups of animals, respectively. Due to the largely different sample sizes of the DOM, BEZ and WIL groups, the resulting total number of alleles scored at each position in the full dataset was also highly dissimilar (1,440 for DOM, 32 for BEZ and 28 for WIL). Considering the different sizes of DOM, BEZ and WIL, we evaluated the effects of applying either a minimum allele frequency (MAF) or a minimum allele count (MAC) threshold. The frequency of the minor allele varied largely in the three groups and precluded the choice of a common threshold for MAF. Applying instead a threshold value based on the Minor Allele Count, not dependent on the number of alleles per group, granted the selection of well documented SNP positions. Considering a maximum sample size of eight, we chose a cutoff value of MAC≥4 corresponding to a frequency of the minor allele of 0.0028 in the DOM group, 0.125 in BEZ and 0.143 in WIL (Figure 1). The MAC≥4 threshold value was then applied on the three subsets of SNPs and the resulting datasets were merged, obtaining a total of 20,695,419 SNPs (20M_BALANCED_MAC dataset) that had MAC≥4 in at least one of the three groups of animals and represented 73.07% of the original dataset. Percentages of SNPs with MAC≥4 in the three clusters and in the merged 20M_BALANCED_MAC dataset indicated that most of the BEZ variants are shared with DOM ones, while the WIL cluster contains a higher proportion of private variants (Figure 2A).

**Figure 1.**
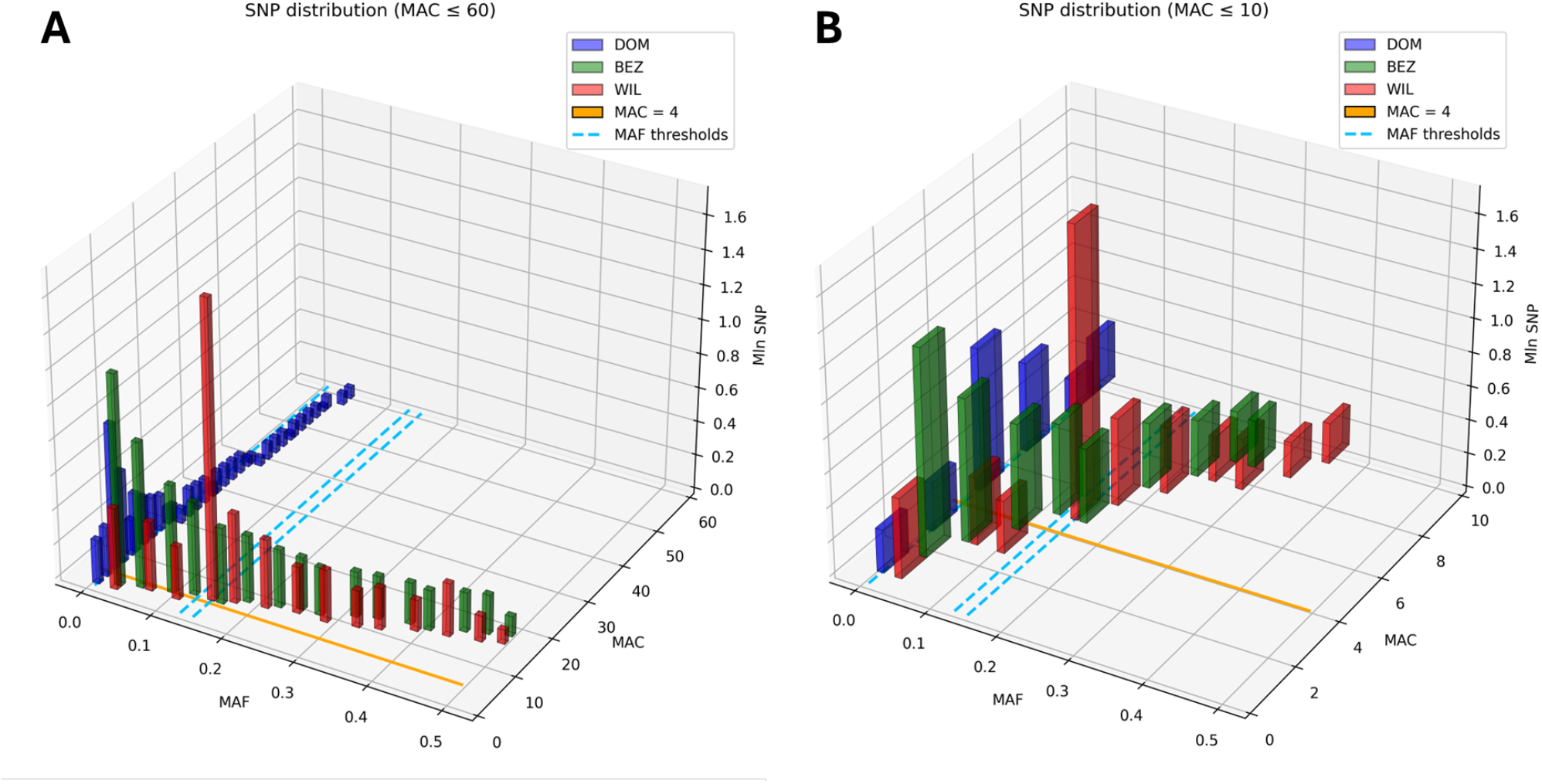
SNP distribution in the domestic, bezoar, and wild datasets. The orange horizontal line (MAC=4) represents the threshold at MAC=4. The light blue dotted lines represent the corresponding thresholds of MAF for DOM (0.0028), BEZ (0.125) and WIL (0.143). SNP were counted within 40 equally spaced bins between MAF=0 and MAF=0.5 or per MAC value from 0 to 60. **Panel A**; SNPs with MAC values from 0 to 60. **Panel B**: SNPs with MAC values from 0 to 10. SNPs with MAC>60 are not shown.

**Figure 2.**
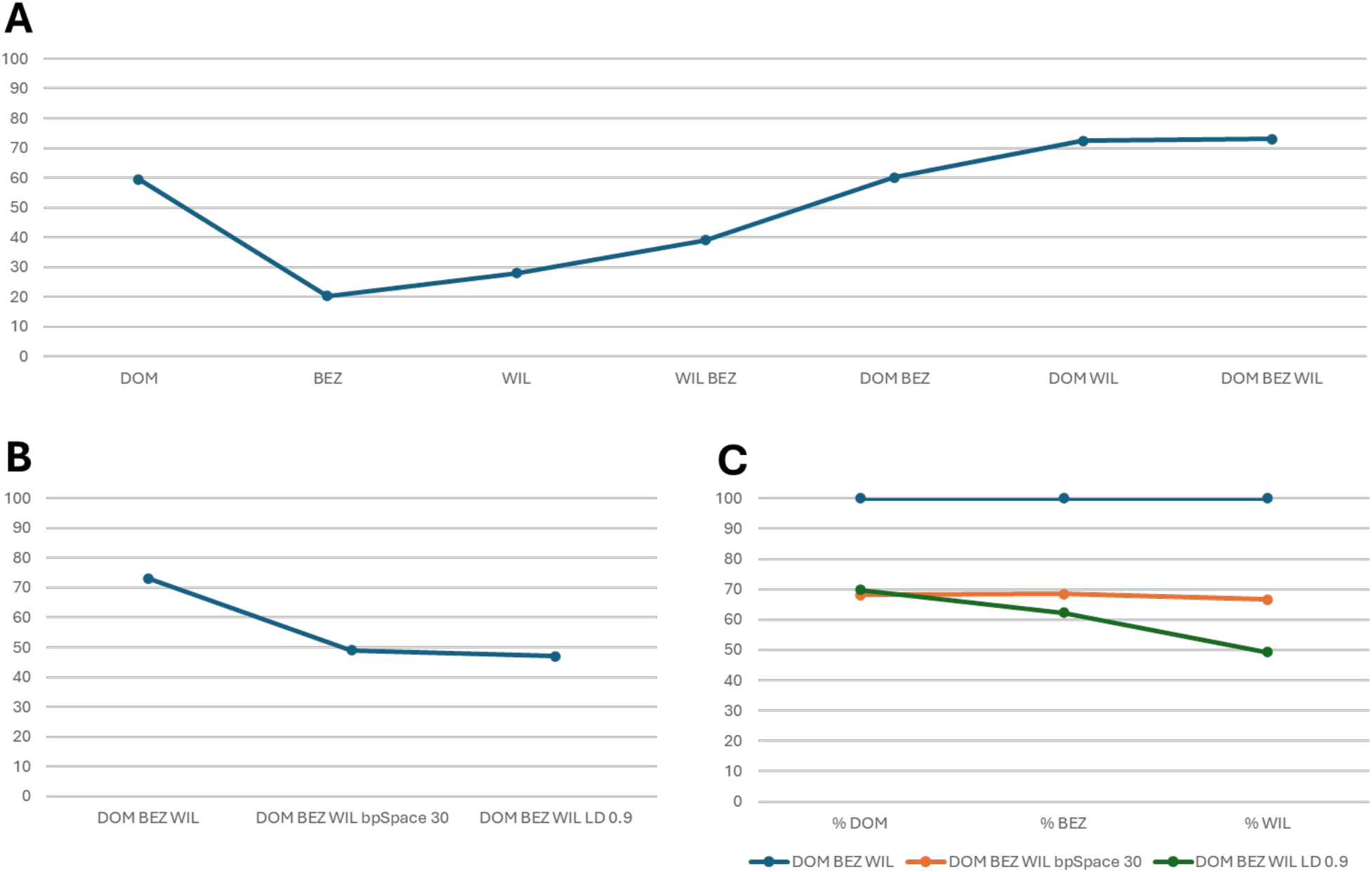
**A**: Percentage of SNPs of the 28M_BALANCED dataset retrieved by filtering the three clusters (DOM, BEZ and WIL) with MAC≥4. Percentages are given for single clusters and merged clusters. **B**: Percentage of SNPs from the 28M_BALANCED dataset surviving filtering with MAC≥4, MAC≥4 + bp-space≥30, and MAC≥4 + LD>0.9 on DOM + BEZ + WIL clusters. **C**: Percentage of cluster-specific SNPs represented in the MAC≥4 + bp-space (orange) and MAC≥4 + LD-filtered (green) datasets, with respect to the MAC≥4 filtered dataset (blue).

### Additional filtering strategy: LD vs. marker spacing

To further thin the dataset, we evaluated the effects of applying additional filtering cutoffs based either on linkage disequilibrium (LD) or on marker spacing to the 20M_BALANCED_MAC dataset. The plink v1.9^22^ bp-space filter excludes one variant from a pair closer than the cutoff distance in bp. The indep-pairwise parameter produces a pruned subset of markers that are in approximate linkage equilibrium with each other. We used a shallow cutoff of LD=0.9 to avoid bias in favor of the most represented gene pools. LD was a little more efficient in thinning the SNP number than bp-space≥30 bps cutoff retaining 47.06 vs 48.96% of the SNPs, respectively (Figure 2B). Compared to bp-space≥30 bps filtering, which retained a percentage for DOM, BEZ and WIL-specific SNPs very similar to the original dataset, LD-based pruning removed a higher number of variants specific for WIL and, to a lesser extent, for BEZ. Therefore, to ensure an even representation of the different gene pools, we adopted the bp-space filtering which returned a dataset of 13,987,126 SNPs (14M_BALANCED dataset) from 750 goats representing 136 breeds and 163 country_breed groups.

In Figure 3, overlaps among the DOM, BEZ and WIL datasets are displayed for the 28M dataset (1372 individuals), as well as for the 14M_BALANCED dataset (750 individuals), showing the proportions of common variants among datasets. Results confirm a higher intersection of variants from domestic goats with those from bezoars, rather than with those from wild goats, and show that the subsampling and filtering procedures did not cause substantial differences in the proportions of variants among species in favour of the most represented.

**Figure 3.**
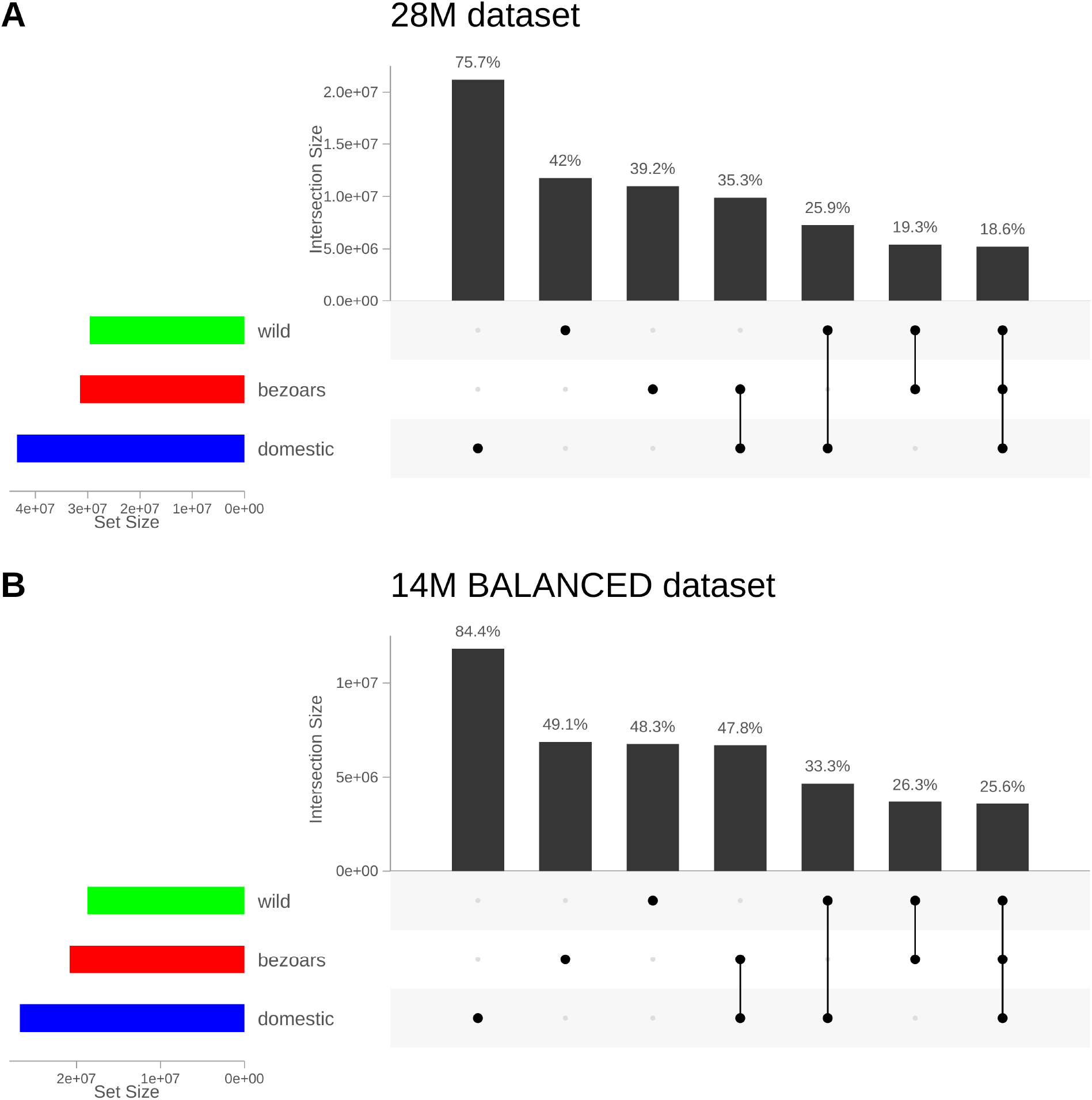
Upset plots of the 28M dataset (Panel A) and of the 14M_BALANCED dataset (Panel B). Numeric intersections of variants across datasets are displayed. Over the bars, the percentage of SNPs with respect to the full dataset is given.

A second dataset focused on domestic goats was prepared using the same parameters and cutoffs starting from the 28M_BALANCED dataset after removing the 30 bezoars and wild goats. This dataset, referred to as the 12M_DOMESTIC dataset, encompassed 11,939,620 SNPs and 720 individuals (128 domestic goat breeds and 149 country_breed groups; Figure 4).

**Figure 4.**
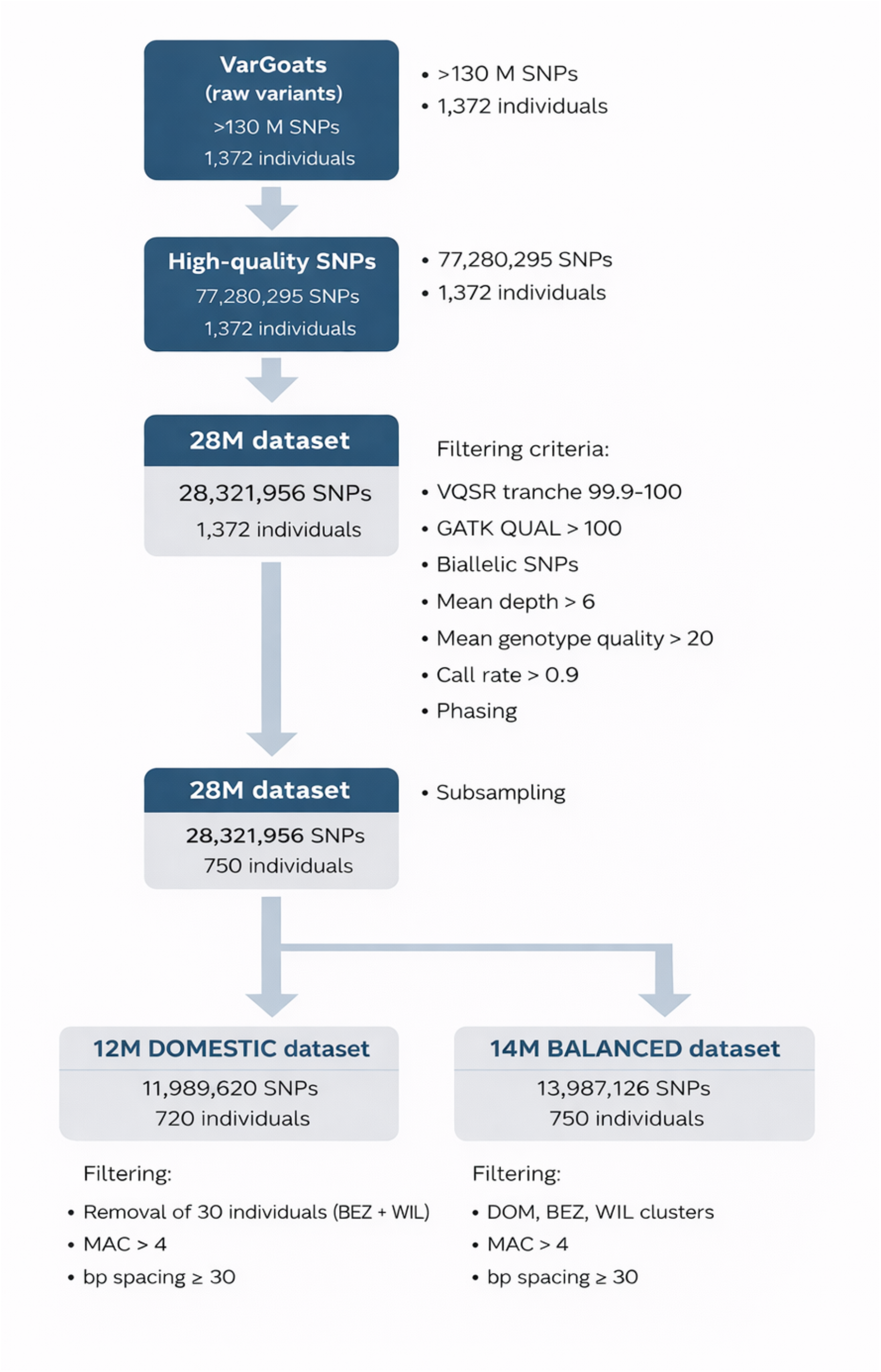
Schematic representation of the filtering procedure applied to the VarGoats dataset.

We verified the effectiveness of the two balanced datasets (750 individuals and 720 individuals) in classifying animals according to species, breed, and geographical provenance, by principal component analysis (PCA), building of neighbor-joining trees (NJ), and model-based clustering by the ADMIXTURE program^23^ on the 12M_DOMESTIC dataset (data partially reported in Figure 5), resulting in a clustering of samples coherent with information contained the available metadata and in line with expectations based on previous results (Colli et al., 2018) and on the known evolutionary history and relationships of goat breeds. To further validate the population structure robustness, additional Admixture runs were performed on 10 subsets, each consisting of 200K randomly extracted SNPs. The plot of cross validation (CV) errors from these runs showed that CV values were highly consistent across runs and indicated the value of K=12 as the best K (Supplementary Figure S1).

**Figure 5.**
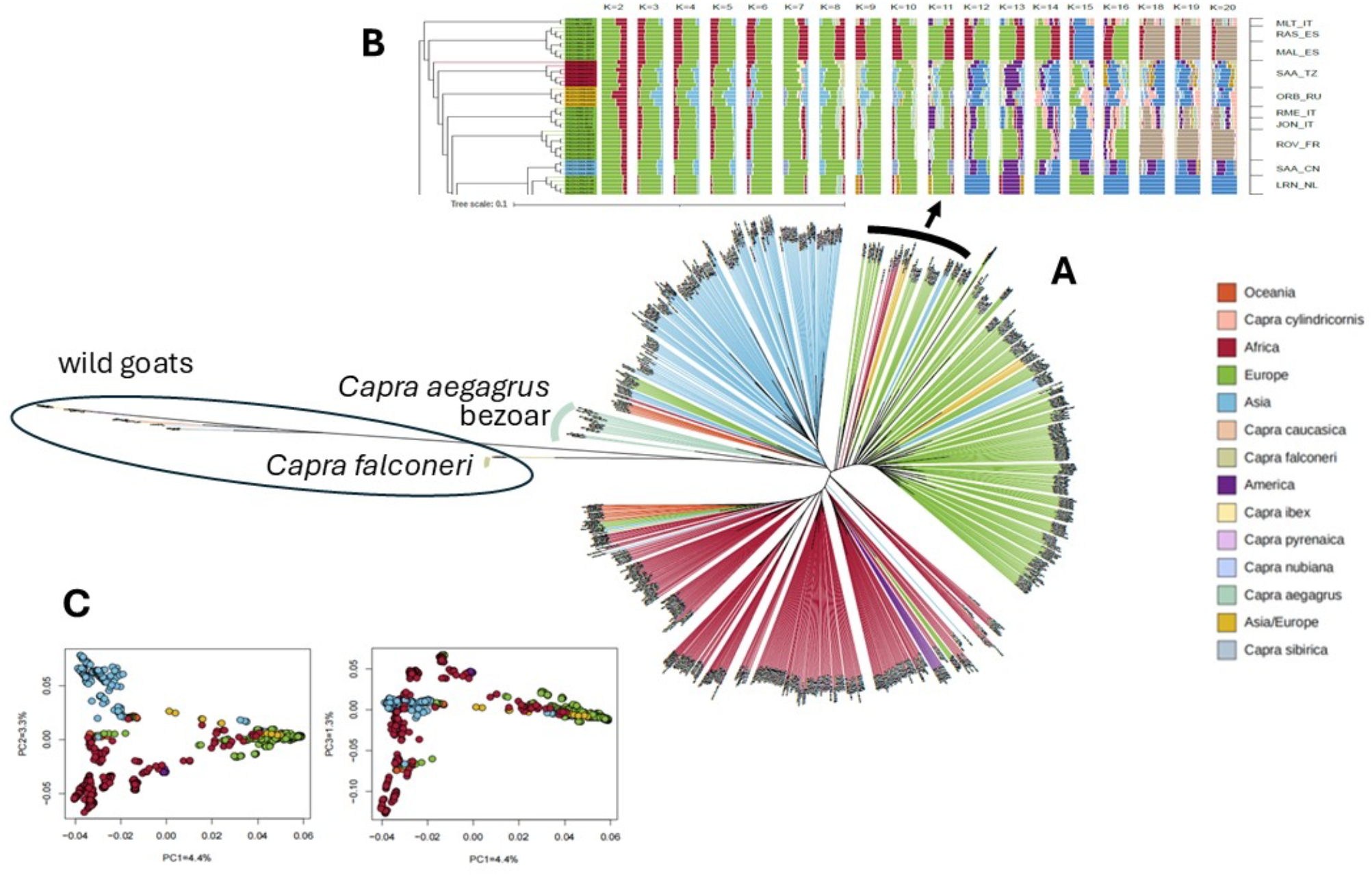
**A**: Neighbor Joining (NJ) tree of the 14M_BALANCED dataset **B**: Admixture results (K2 to K20) for an example subset of VarGoats animals, showing clustering of Saanen country_breeds sampled outside Europe within the European cluster (SAA_TZ = Saanen_Tanzania, ORB_RU = Orenburg Cachemire_Russia, SAA_CN = Saanen_China). **C**: Principal Component analysis of the 12M_DOMESTIC dataset.

The NJ tree of the 750 individuals 14M_BALANCED dataset (Figure 5A) displayed a good resolution of wild goat species, and also resolved domestic goats according to their continental provenance. Populations of transboundary or cosmopolitan breeds were correctly assigned to their ancestral gene pool of origin (Figure 5B). The PCA carried out on the 720 individuals 12M_DOMESTIC dataset (Figure 5C) confirmed Admixture and NJ tree results obtained before.

## Conclusions

Due to the decreasing cost of resequencing data, the last few years have seen an increase in the production of genomic data for many species, including domestic ones. In most cases, the datasets that can be assembled starting from these data are often highly unbalanced in terms of representativeness of local gene pools’ variation, with some populations or breeds being overrepresented. Also, these genomic collections are usually large-sized and require big computational facilities to be analyzed, thus preventing research groups with limited computation resources from being autonomous in driving data analyses. Therefore, it is necessary to proceed with variant filtering before processing data in diversity studies, but strategies that allow to conserve the representative fraction of the total variation must be applied. The VarGoats dataset is an example of these occurrences since it represents a collection of whole-genome sequences that were produced based on requirements of different projects, and are therefore highly variable in terms of number of animals per breed, geographical provenance, farming systems, year of sampling and other recorded metadata. The subsampling and filtering procedures adopted to thin the VarGoats 28M dataset to a more manageable 14M/12M set proved to be adequate to retain the information content mirroring the gene pools of wild goat species, of both transboundary and local breeds and of local gene pools characterizing different geographical areas. These sets of markers are suitable for further goat diversity analyses aiming at e.g. the reconstruction of evolutionary history of goat populations across the world, the identification of signatures of selection, or the study of the genetic bases of environmental adaptation. Other analyses, i.e. genetic distances, that require much smaller datasets or unlinked markers could benefit from further filtering based on different parameters. Thus, according to the kind of analysis to be performed, additional filtering steps can be advisable.

The strategies and parameters adopted in this study can be applied to datasets with features like VarGoats, when problems like unequal distribution of individuals across breeds and species are encountered and a reduction in marker numbers is desirable. In this work, cutoffs were set as not to be aggressive in drastically reducing the number of variants, but to decrease redundancy without losing the information contained in the dataset and without introducing biases caused by its heterogeneity. Nonetheless, thresholds can be adjusted according to the target dataset size, provided that a careful inspection of the reduced dataset is carried out to verify that its original information content has been retained and no bias has been introduced during the filtering steps.

## Materials and methods

### Variant quality filtering

The entire dataset comprises animals from the VarGoats collection, which were sequenced at 15x coverage on an Illumina HiSeq X sequencer^19^, and animals from public databases. In Table 1 and Supplementary Table S1, individuals included in the dataset are reported, according to their distribution across breeds and species. The 1,372 animals were mapped to the ARS1 *C. hircus* genome (GCA_001704415.1)^20^ with the BWA-MEM software (v0.7.X). Variants were called with GATK-HaplotypeCaller^24^, then, preliminary quality filters were applied using VQSR (keeping the 99.9 to 100.00 tranche), selecting variants with GATK quality >100 and retaining only biallelic SNPs, resulting in a dataset of 77,280,295 SNPs^19^. Outlier animals were identified through the analysis of a subset of SNPs included in the Goat_IGGC_65K_v2 array (https://www.goatgenome.org/projects.html#50K_snp_chip2) extracted for the VCF file, after quality control on the markers. Observed heterozygosity (H_O_), PCA and Admixture were used to detect outlier individuals. For the H_O_ for each breed the individuals with a level of H_O_ above Q3 + 1.5xIQR or below Q1 - 1.5xIQR were considered outliers and removed from the dataset before performing further quality filtering steps on VCF files. Subsequent simulations with different cutoffs were run to evaluate filtering effects on the number of surviving variants (Supplementary Figure S2) and on breed representation in the dataset. The mean depth (DP) >6, mean genotype quality (GQ) >20, and call rate (CR) >0.9 filters were applied with BCFtools^25^, and 28,645,747 SNPs were retrieved. Outliers that had previously been removed for filtering cutoffs definition were newly added to the dataset. Most of them were considered mislabeled and taken into consideration for relabeling. Phasing was then performed with Beagle 5.3 with standard parameters, and unphased SNPs were removed. The resulting dataset encompasses 28,321,956 quality-filtered, biallelic, phased SNPs (28M dataset).

### Relabeling and subsampling

The identification of misbehaving samples and their eventual relabeling were based on the evaluation of genotype data at a subset of autosomal SNPs from the Illumina GoatSNP60 BeadChip^26^. A subset of 59,929 SNPs, present in chromosomes 1-29 and MT in the Illumina 60K, was extracted from the VCF files and compared with the same markers typed with the 60K array over the same individuals. The resulting raw datasets were both filtered by MAF <0.01 and CR >0.10. Animals with concordance rate between extracted SNPs and SNP chip data below 70% were marked and considered mislabeled in one of the two datasets. The first 5 principal components (PCs, about 15% of explained variance in total) were used in PCA analysis, and animals were marked as outliers in each PC if exceeding the 1^st^ or 3^rd^ quartile of more than IQR*1.5 (Inter Quartile Range, IQR). Animals classified as outliers in more than 60% of the cases were marked. An exploratory population structure analysis was carried out with ADMIXTURE software^23^ with a fivefold Cross-Validation error calculation, based on which K=8 was identified as the best fitting solution. In each of the eight clusters, animals were identified as outliers if exceeding the 1^st^ or 3^rd^ quartile of more than IQR*1.5. Animals classified as outliers in more than 60% of the cases were marked. Observed Heterozygosity H_O_ was also calculated, finding a mean value of H_O_= 0.3477. Animals were marked if their H_O_ was higher than 0.5392 or lower than 0.1561. Other clustering approaches as Neighbor-joining trees based on Allele Sharing Distance (ASD) were further used to check animals’ classification and mark as outlier individuals not clustering within their declared breed. When possible, animals were relabeled with the correct breed assignment according to the results obtained with the described evaluations, otherwise they were marked as unknown breed (UNK). A subsampling procedure was then performed to exclude subjects with problems related either to the quality of the genomic data or to the behavior with respect to the assigned breed in any of the control analyses, resulting in the removal of 15 animals from 6 underrepresented *C. hircus* country_breeds.

After the subsampling step based on individual quality and breed assignment, BITE v2.0^21^ was used to select animals to be retained within those belonging to overrepresented country_breeds, setting at 8 the maximum number of animals per country_breed.

### Thinning procedures and filtered datasets validation

The analyses required to select and test parameters to be applied for variant filtering were performed with plink v1.9^22^ (--maf, --mac, --bp-space and --indep-pairwise parameters), VCFtools^25^, BCFtools and custom scripts.

The GCTA software^27^ was used to estimate the genetic relationship matrix (GRM) between pairs of individuals from a set of SNPs and perform the PCA analysis. A distant matrix was computed with plink v1.9 and used to prepare the Neighbor Joining tree with MEGA^28^. The software ADMIXTURE was run with default parameters and five-fold cross-validation for K from 2 to 8.

## Data availability

VarGoats sequence datasets are available at the NCBI BioProject db, accession PRJEB74076^29^, PRJEB74075^30^, PRJEB50463^31^, PRJEB37507^32^, PRJEB37841^33^, PRJEB37276^34^, PRJEB37122^35^, PRJEB37208^36^, PRJEB31857^37^, PRJNA529091^38^, PRJEB3134^39^, PRJEB3135^40^, PRJEB3136^41^, PRJEB4371^42^, PRJEB5166^43^, and PRJEB5900^44^. Variants are accessible on the European Variation Archive under accession PRJEB90141.

Use of these data is regulated by a data sharing agreement which is available here: http://www.goatgenome.org/vargoats_agreement.html.

## Acknowledgements

The authors wish to thank Dr. Benjamin D. Rosen and Dr. Mahesh Neupane from the Animal Genomics and Improvement Laboratory, USDA-ARS, for phasing the 28M SNP dataset and Dr. Bertrand Servin and Dr. Pierre Faux (GenPhySE, INRAE) for the exchange of opinions regarding the treatment of the data. AT is funded by the Jamieson Bequest fund at the School of Biodiversity, One-Health and Veterinary Medicine of the University of Glasgow.

The VarGoats project members: Alessandra Crisà (Council for Agricultural Research and Economics CREA, Research Centre for Animal Production and Aquaculture, Monterotondo, Italy); Andrea Talenti (University of Glasgow, School of Biodiversity, One-Health and Veterinary Medicine, Glasgow, United Kingdom); Antonia Noce (Universitat Autònoma de Barcelona, CRAG Centre de Recerca en Agrigenòmica, Barcelona, Spain); Arianna Bionda (University of Milan, Dipartimento di Scienze Agrarie e Ambientali DiSAA, Milan, Italy); Arianna Manunza (Georg-August Universität Göttingen, Fakultät für Forstwissenschaften und Waldökologie, Abteilung Wildtierwissenschaften, Göttingen, Germany); Badr Benjelloun (INRAE-Moroc, Rabat, Morocco); Barbara Lazzari (National Research Council of Italy CNR, IBBA Istituto di Biologia e Biotecnologia Agraria, Milan, Italy); Ben Rosen (USDA, Animal Genomics and Improvement Laboratory, Beltsville, MD, USA); Bertrand Servin (INRAE, Génétique Physiologie et Systèmes d’Elevage GenPhySE, Castanet Tolosan, France); Carine Genet (INRAE, Génétique Physiologie et Systèmes d’Elevage GenPhySE, Castanet Tolosan, France); Carole Charlier (University of Liège, GIGA, Liège, Belgium); Caroline Leroux (Université Lyon 1, INRAE, Lyon, France); Coralie Danchin (IDELE, Génétique et gestion des populations animales, Paris, France); Cord Drögemüller (University of Bern, Institute of Genetics, Bern, Switzerland); Curt Van Tassell (USDA Animal Genomics and Improvement Laboratory, Beltsville, MD, USA); Diego Alonso Vargas Donayre (Universitat Autònoma de Barcelona, CRAG Centre de Recerca en Agrigenòmica, Barcelona, Spain); Elena Petretto (Università Cattolica del S. Cuore, Department of Animal Science, Food and Nutrition DIANA, Piacenza, Italy); Emilio Mármol-Sánchez (Center for Evolutionary Hologenomics, The Globe Institute, University of Copenhagen, Copenhagen, Denmark); Emily Clark (EMBL-EBI European Bioinformatics Institute, Hinxton, United Kingdom); Emmanuelle Lerat (Université Lyon 1, CNRS Biometrie et Biologie Evolutive, Villeurbanne, France); François Pompanon (Université Grenoble Alpes, Laboratoire d’Ecologie Alpine, Grenoble, France); George Liu (USDA Animal Genomics and Improvement Laboratory, Beltsville, MD, USA); Gwenola Tosser-Klopp (INRAE GenPhySE, Castanet Tolosan, France); Imen Baazaoui (Universitat Autònoma de Barcelona, CRAG Centre de Recerca en Agrigenòmica, Barcelona, Spain); Johannes Arjen Lenstra (Utrecht University, Faculty of Veterinary Medicine, Utrecht, Netherlands); Joram M. Mwacharo (International Centre for Agricultural Research in the Dry Areas ICARDA, Addis Ababa, Ethiopia); James Prendergast (University of Edinburgh, Roslin Institute, Midlothian, United Kingdom); Lin Jiang (Chinese Academy of Agricultural Sciences, Institute of Animal Science, Beijing, China); Jocelyn Turpin (Université Lyon 1, INRAE, Lyon, France); Jolijn Erven (University College Dublin, School of Agriculture and Food Science, Dublin, Ireland); Kevin Daly (University College Dublin, School of Agriculture and Food Science, Dublin, Ireland); Laura Botigué (Universitat Autònoma de Barcelona, CRAG Centre de Recerca en Agrigenòmica, Barcelona, Spain); Laurence Drouilhet (INRAE GenPhySE, Castanet Tolosan, France); Licia Colli (Università Cattolica del S. Cuore, Department of Animal Science, Food and Nutrition DIANA, Piacenza, Italy); Lingzhao Fang (University of Edinburgh, Roslin Institute, Midlothian, United Kingdom); Marina Naval (CSIRO Livestock genomics, St. Lucia, Queensland, Australia); Mahesh Neupane (USDA Animal Genomics and Improvement Laboratory, Beltsville, MD, USA); Marcel Amills (Universitat Autònoma de Barcelona, CRAG Centre de Recerca en Agrigenòmica, Barcelona, Spain); Marco Milanesi (Università Cattolica del S. Cuore, Department of Animal Science, Food and Nutrition DIANA, Piacenza, Italy); Maria Gracia Luigi Sierra (Universitat Autònoma de Barcelona, CRAG Centre de Recerca en Agrigenòmica, Barcelona, Spain); Mario Barbato (Università degli Studi di Messina, Dipartimento di Scienze Veterinarie, Messina, Italy); Matteo Cortellari (University of Milan, Dipartimento di Scienze Agrarie e Ambientali DiSAA, Milan, Italy); Maxime Ben Braiek (INRAE, Génétique Physiologie et Systèmes d’Elevage GenPhySE, Castanet Tolosan, France); Mazdak Salavati (Scotland’s Rural College SRUC, Dairy Research and Innovation Centre, Edinburgh, United Kingdom); Mingjing Wang (Universitat Autònoma de Barcelona, CRAG Centre de Recerca en Agrigenòmica, Barcelona, Spain); Paola Crepaldi (University of Milan, Dipartimento di Scienze Agrarie e Ambientali DiSAA, Milan, Italy); Philippe Bardou(INR AE, Génétique Physiologie et Systèmes d’Elevage GenPhySE, Castanet Tolosan, France); Pierre Faux (INRAE, Génétique Physiologie et Systèmes d’Elevage GenPhySE, Castanet Tolosan, France); Rachel Rupp (INRAE, Génétique Physiologie et Systèmes d’Elevage GenPhySE, Castanet Tolosan, France); Roberto Steri (Council for Agricultural Research and Economics CREA, Research Centre for Animal Production and Aquaculture, Monterotondo, Italy); Rudiger Brauning (AgResearch ltd., Bioeconomy Science Institute, Mosgiel, New Zealand); Alessandra Stella (National Research Council of Italy CNR, IBBA Istituto di Biologia e Biotecnologia Agraria, Milan, Italy); Stéphane Fabre (INRAE, Génétique Physiologie et Systèmes d’Elevage GenPhySE, Castanet Tolosan, France); Taina Figueiredo (Universitat Autònoma de Barcelona, CRAG Centre de Recerca en Agrigenòmica, Barcelona, Spain); Thomas Faraut (INRAE, Génétique Physiologie et Systèmes d’Elevage GenPhySE, Castanet Tolosan, France); Valentin Sorin (INRAE, GABI and GenPHYse, Jouy-en-Josas, France); Ke Wang (Universitat Autònoma de Barcelona, CRAG Centre de Recerca en Agrigenòmica, Barcelona, Spain); Clet Wandui Masiga (Tropical Institute of Development Innovations TRIDI, Kampala, Uganda); Wilson Nandolo (Lilongwe University of Agriculture and Natural Resources, Bunda Animal Breeding Centre, Lilongwe, Malawi); Yefang Li (Chinese Academy of Agricultural Sciences, Institute of Animal Science, Beijing, China); Zexi Cai (Aarhus university, Center for Quantitative Genetics and Genomics, Aarhus, Denmark).

## Author contributions

BL, PC, LC, and GTK developed the research idea and sampling design and acquired funding; AB and JAL contributed to the relabeling and subsampling procedures; BL, YL, MM, PB, LJ and AT performed the data analyses; BL, MM, AT, AB, PC and LC performed the data interpretation. BL and LC wrote the paper. All authors edited and approved the final manuscript.

## Ethics declarations

### Competing interests

The authors declare that they have no competing interests.

## Notes

### Competing Interest Statement

The authors have declared no competing interest.

### Summary of Updates

This version of the manuscript has been revised to include the VarGoats Consortium in the Authors list and the full list of the VarGoats members in the Acknowledgements section. Figures quality was improved and small changes were made to the text.

