## Supplemental Table and Figures for "A tailored variant filtering procedure for multi-breed and multi-species unbalanced animal SNP collections"

**Supplementary Table S1.** Composition of the analysed dataset.

| Breed | Species | Nr of animals | Country | Continental area |
|---|---|---|---|---|
| Abadah | *Capra hircus* | 1 | Iran | Asia |
| Abergelle | *Capra hircus* | 10 | Ethiopia | Africa |
| Alpine | *Capra hircus* | 58 | France | Europe |
| Alpine | *Capra hircus* | 12 | Italy | Europe |
| Alpine | *Capra hircus* | 22 | Switzerland | Europe |
| Androy | *Capra hircus* | 5 | Madagascar | Africa |
| Angora | *Capra hircus* | 20 | France | Europe |
| Angora | *Capra hircus* | 6 | Madagascar | Africa |
| Angora | *Capra hircus* | 5 | Pakistan | Asia |
| Angora | *Capra hircus* | 3 | South Africa | Africa |
| Anhui White | *Capra hircus* | 1 | China | Asia |
| Appenzell | *Capra hircus* | 24 | Switzerland | Europe |
| Balaka Ulongwe | *Capra hircus* | 4 | Malawi | Africa |
| Bange Cashmere | *Capra hircus* | 1 | China | Asia |
| Barbari | *Capra hircus* | 1 | Pakistan | Asia |
| Bari | *Capra hircus* | 5 | Pakistan | Asia |
| Beetal | *Capra hircus* | 10 | Pakistan | Asia |
| Bengal | *Capra hircus* | 7 | Bangladesh | Asia |
| Bermeya | *Capra hircus* | 6 | Spain | Europe |
| Boer | *Capra hircus* | 7 | Australia | Oceania |
| Boer | *Capra hircus* | 4 | Korea | Asia |
| Boer | *Capra hircus* | 8 | New Zealand | Oceania |
| Boer | *Capra hircus* | 5 | Switzerland | Europe |
| Boer | *Capra hircus* | 3 | Tanzania | Africa |
| Boer | *Capra hircus* | 8 | Zimbabwe | Africa |
| Boiragi | *Capra hircus* | 1 | Bangladesh | Asia |
| Burundi | *Capra hircus* | 2 | Switzerland | Europe |
| Cashmere | *Capra hircus* | 7 | Australia | Oceania |
| Cashmere | *Capra hircus* | 15 | China | Asia |
| Cashmere | *Capra hircus* | 1 | India | Asia |
| Chaidamu | *Capra hircus* | 1 | China | Asia |
| Chengde Hornless | *Capra hircus* | 1 | China | Asia |
| Chengdu Grey | *Capra hircus* | 1 | China | Asia |
| Ciociara Grigia | *Capra hircus* | 4 | Italy | Europe |
| Creole | *Capra hircus* | 16 | Guadeloupe | America |
| Damani | *Capra hircus* | 5 | Pakistan | Asia |
| Danish Landrace | *Capra hircus* | 6 | Denmark | Europe |
| Dedza | *Capra hircus* | 4 | Malawi | Africa |
| Deradin Panah | *Capra hircus* | 5 | Pakistan | Asia |
| Diana | *Capra hircus* | 8 | Madagascar | Africa |
| Djallonke | *Capra hircus* | 1 | Burkina Faso | Africa |
| Dutch Landrace | *Capra hircus* | 5 | Netherlands | Europe |
| Finnish Landrace | *Capra hircus* | 6 | Finland | Europe |
| Fosses | *Capra hircus* | 16 | France | Europe |
| Galla | *Capra hircus* | 8 | Kenya | Africa |

| | | | | |
|---|---|---|---|---|
| Gaza | *Capra hircus* | 4 | Mozambique | Africa |
| Girgentana | *Capra hircus* | 4 | Italy | Europe |
| Gogo | *Capra hircus* | 8 | Tanzania | Africa |
| Grisons Striped | *Capra hircus* | 25 | Switzerland | Europe |
| Guera | *Capra hircus* | 8 | Mali | Africa |
| Guishan Black | *Capra hircus* | 1 | China | Asia |
| Guizhou Black | *Capra hircus* | 1 | China | Asia |
| Gumez | *Capra hircus* | 8 | Ethiopia | Africa |
| Icelandic | *Capra hircus* | 5 | Iceland | Europe |
| Iran North Central | *Capra hircus* | 22 | Iran | Asia |
| Iran South | *Capra hircus* | 3 | Iran | Asia |
| Iraq | *Capra hircus* | 1 | Iraq | Asia |
| Jianchang Black | *Capra hircus* | 1 | China | Asia |
| Jining Grey | *Capra hircus* | 41 | China | Asia |
| Jonica | *Capra hircus* | 4 | Italy | Europe |
| Kachan | *Capra hircus* | 5 | Pakistan | Asia |
| Kamori | *Capra hircus* | 5 | Pakistan | Asia |
| Keffa | *Capra hircus* | 8 | Ethiopia | Africa |
| Khalkhali | *Capra hircus* | 1 | Iran | Asia |
| Khurasani | *Capra hircus* | 5 | Pakistan | Asia |
| Korea Native Goat | *Capra hircus* | 27 | Korea | Asia |
| Kurdi | *Capra hircus* | 2 | Iran | Asia |
| Lai Wublack | *Capra hircus* | 1 | China | Asia |
| Landin | *Capra hircus* | 10 | Mozambique | Africa |
| Laoshan | *Capra hircus* | 2 | China | Asia |
| Leizhou | *Capra hircus* | 6 | China | Asia |
| Liaoning Cashmere | *Capra hircus* | 5 | China | Asia |
| Lilongwe | *Capra hircus* | 3 | Malawi | Africa |
| Longlin | *Capra hircus* | 6 | China | Asia |
| Lorraine | *Capra hircus* | 13 | France | Europe |
| Lvliang Black | *Capra hircus* | 1 | China | Asia |
| Maasai | *Capra hircus* | 8 | Tanzania | Africa |
| Maguan Horn Down | *Capra hircus* | 1 | China | Asia |
| Mallorquina | *Capra hircus* | 6 | Spain | Europe |
| Maltese | *Capra hircus* | 2 | Italy | Europe |
| Malya | *Capra hircus* | 8 | Tanzania | Africa |
| Manica | *Capra hircus* | 3 | Mozambique | Africa |
| Mashona | *Capra hircus* | 12 | Zimbabwe | Africa |
| Matebele | *Capra hircus* | 8 | Zimbabwe | Africa |
| Matou | *Capra hircus* | 1 | China | Asia |
| Maure | *Capra hircus* | 4 | Mali | Africa |
| Menabe | *Capra hircus* | 8 | Madagascar | Africa |
| Montecristo | *Capra hircus* | 5 | Italy | Europe |
| Morocco | *Capra hircus* | 163 | Morocco | Africa |
| Mubende | *Capra hircus* | 3 | Uganda | Africa |
| Nachi | *Capra hircus* | 5 | Pakistan | Asia |
| Nadjdi | *Capra hircus* | 1 | Iran | Asia |
| Naine | *Capra hircus* | 8 | Mali | Africa |

| | | | | |
|---|---|---|---|---|
| Nera Verzasca | *Capra hircus* | 24 | Switzerland | Europe |
| Nsanje | *Capra hircus* | 6 | Malawi | Africa |
| Old Irish Goat | *Capra hircus* | 5 | Ireland | Europe |
| Orenburg Cashmere | *Capra hircus* | 5 | Russia | Asia/Europe |
| Pafuri | *Capra hircus* | 3 | Mozambique | Africa |
| Palmera | *Capra hircus* | 6 | Spain | Europe |
| Pare White | *Capra hircus* | 8 | Tanzania | Africa |
| Pateri | *Capra hircus* | 4 | Pakistan | Asia |
| Peacock | *Capra hircus* | 24 | Switzerland | Europe |
| Peulh | *Capra hircus* | 4 | Mali | Africa |
| Poitevine | *Capra hircus* | 12 | France | Europe |
| Provencale | *Capra hircus* | 15 | France | Europe |
| Pyrenean | *Capra hircus* | 15 | France | Europe |
| Qinghai | *Capra hircus* | 6 | China | Asia |
| Raini | *Capra hircus* | 1 | Iran | Asia |
| Rangeland | *Capra hircus* | 2 | Australia | Oceania |
| Raoshan White | *Capra hircus* | 1 | China | Asia |
| Rasquera | *Capra hircus* | 4 | Spain | Europe |
| Ritu Cashmere | *Capra hircus* | 1 | China | Asia |
| Rossa Mediterranea | *Capra hircus* | 3 | Italy | Europe |
| Rove | *Capra hircus* | 11 | France | Europe |
| Saanen | *Capra hircus* | 5 | China | Asia |
| Saanen | *Capra hircus* | 36 | France | Europe |
| Saanen | *Capra hircus* | 5 | Italy | Europe |
| Saanen | *Capra hircus* | 10 | Korea | Asia |
| Saanen | *Capra hircus* | 6 | Russia | Asia/Europe |
| Saanen | *Capra hircus* | 23 | Switzerland | Europe |
| Saanen | *Capra hircus* | 8 | Tanzania | Africa |
| Savoie | *Capra hircus* | 15 | France | Europe |
| Shaanan White | *Capra hircus* | 1 | China | Asia |
| Sichuan | *Capra hircus* | 1 | China | Asia |
| Small East African | *Capra hircus* | 7 | Kenya | Africa |
| Small East African | *Capra hircus* | 7 | Mozambique | Africa |
| Sofia | *Capra hircus* | 8 | Madagascar | Africa |
| Sonjo | *Capra hircus* | 8 | Tanzania | Africa |
| Soudanaise | *Capra hircus* | 8 | Mali | Africa |
| St Gallen Booted | *Capra hircus* | 24 | Switzerland | Europe |
| Sud Ouest | *Capra hircus* | 8 | Madagascar | Africa |
| Tali | *Capra hircus* | 1 | Iran | Asia |
| Tanzanian Norwegian | *Capra hircus* | 7 | Tanzania | Africa |
| Targui | *Capra hircus* | 4 | Mali | Africa |
| Teddy | *Capra hircus* | 5 | Pakistan | Asia |
| Tessin Grey | *Capra hircus* | 13 | Switzerland | Europe |
| Thari | *Capra hircus* | 5 | Pakistan | Asia |
| Thyolo | *Capra hircus* | 8 | Malawi | Africa |
| Tibetan | *Capra hircus* | 25 | China | Asia |
| Toggenburg | *Capra hircus* | 1 | Kenya | Africa |
| Toggenburg | *Capra hircus* | 24 | Switzerland | Europe |

| | | | | |
|---|---|---|---|---|
| Toggenburg | *Capra hircus* | 8 | Tanzania | Africa |
| Tunisian | *Capra hircus* | 5 | Tunisia | Africa |
| Turki Qashqai | *Capra hircus* | 1 | Iran | Asia |
| Unknown | *Capra hircus* | 1 | Australia | Oceania |
| Unknown | *Capra hircus* | 4 | Switzerland | Europe |
| Valais | *Capra hircus* | 27 | Switzerland | Europe |
| Valais Copper Neck | *Capra hircus* | 1 | Switzerland | Europe |
| Valdostana | *Capra hircus* | 2 | Italy | Europe |
| Woyito Guji | *Capra hircus* | 8 | Ethiopia | Africa |
| Wu Zhu Mu Qin White | *Capra hircus* | 1 | China | Asia |
| Xiang Dong Black | *Capra hircus* | 1 | China | Asia |
| Xinjiang | *Capra hircus* | 2 | China | Asia |
| Yimeng | *Capra hircus* | 1 | China | Asia |
| Zhon Wei | *Capra hircus* | 1 | China | Asia |
| Bezoar | *Capra aegagrus* | 20 | Iran | Asia |
| Bezoar | *Capra aegagrus* | 10 | Turkey | Asia |
| West Caucasian Tur | *Capra caucasica* | 1 | France | Europe |
| West Caucasian Tur | *Capra caucasica* | 1 | Russia | Asia |
| East Caucasian Tur | *Capra cylindricornis* | 1 | Azerbaijan | Asia |
| Markhor | *Capra falconeri* | 1 | France | Europe |
| Markhor | *Capra falconeri* | 1 | Uzbekistan | Asia |
| Alpine Ibex | *Capra ibex* | 1 | France | Europe |
| Alpine Ibex | *Capra ibex* | 4 | Italy | Europe |
| Alpine Ibex | *Capra ibex* | 1 | Unknown | Europe |
| Nubian Ibex | *Capra nubiana* | 1 | Israel | Asia |
| Pyrenean Ibex | *Capra pyrenaica* | 1 | France | Europe |
| Siberian Ibex | *Capra sibirica* | 1 | Pakistan | Asia |
| Siberian Ibex | *Capra sibirica* | 1 | Tajikistan | Asia |

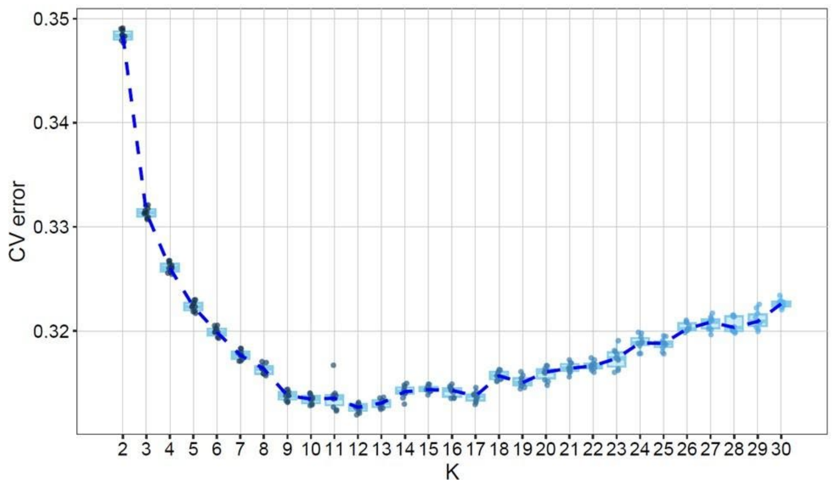


**Supplementary Figure S1**

Admixture CV error plot of 10 subsets, each consisting of 200K SNPs randomly extracted from the 720 individuals balanced dataset (only domestic goats).

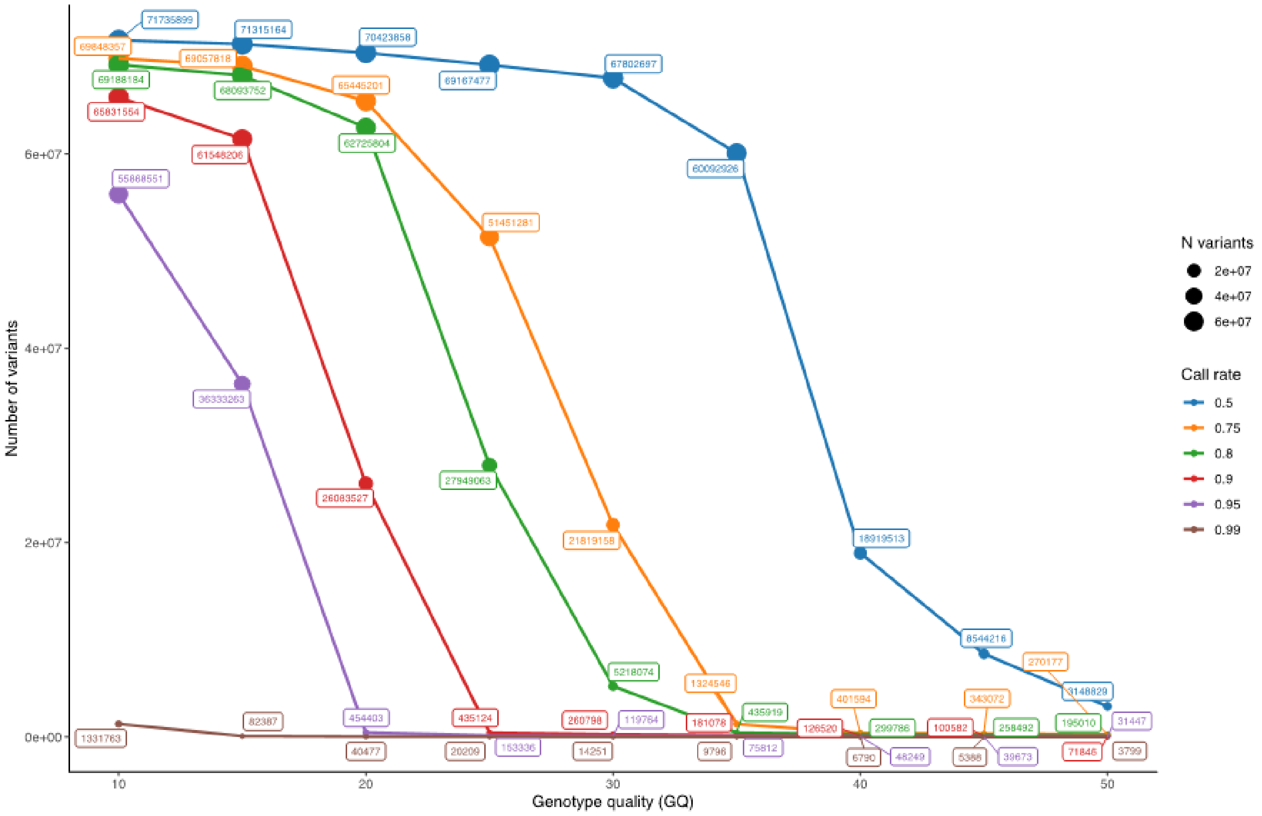


### Supplementary Figure S2

Plot of the effects of filtering according to genotype quality (GQ) and call rate on the number of surviving variants.
